# Co-occurrences enhance our understanding of aquatic fungal metacommunity assembly and reveal potential host–parasite interactions

**DOI:** 10.1101/2022.06.21.496979

**Authors:** Máté Vass, Karolina Eriksson, Ulla Carlsson-Graner, Johan Wikner, Agneta Andersson

## Abstract

Our knowledge of aquatic fungal communities, their assembly, distributions and ecological roles in marine ecosystems is scarce. Hence, we aimed to investigate fungal metacommunities of coastal habitats in a subarctic zone (northern Baltic Sea, Sweden). Using a novel joint species distribution model and network approach, we quantified the importance of biotic associations contributing to the assembly of mycoplankton, further, detected potential biotic interactions between fungi–algae pairs, respectively. Our long-read metabarcoding approach identified 504 fungal taxa, of which a dominant fraction (44.8 %) was assigned as early-diverging fungi (i.e., Cryptomycota and Chytridiomycota). Alpha diversity of mycoplankton declined and community compositions changed along inlet–bay– offshore transects. The distributions of most fungi were rather influenced by spatial factors than by environmental drivers, and the influence of biotic associations was pronounced when environmental filtering was weak and spatial patterning lessened. We found great number of co-occurrences (138) among the dominant fungal groups, and the forty associations between fungal and algal OTUs suggested potential host–parasite/saprotroph links, supporting a Cryptomycota-based mycoloop pathway. We emphasize that the contribution of biotic associations to mycoplankton assembly are important to consider in future studies as it helps to improve predictions of species distributions in aquatic ecosystems.

## Introduction

Aquatic fungi comprise a diverse group of heterotrophic microorganisms, spanning a wide range of life strategies from saprotrophy through mutualism to parasitism (Nilsson et al. 2019). Their contribution and impact on the biogeochemical processes of the biosphere is crucial (Grossart & Rojas-Jimenez 2016, Gladfelter et al. 2019, Grossart et al. 2019).

Molecular analysis of environmental samples using next-generation sequencing have unravelled an extremely high diversity of undescribed fungi (see e.g., Richards et al. 2015) that is often referred as the “dark matter fungi”, or DMF (Grossart et al. 2015). DMF are commonly found within the early-diverging lineages of the fungal tree of life, including members of the basal phyla of Cryptomycota and Chytridiomycota. These fungi have shown to be saprotrophs and obligate or facultative parasites, however, very limited knowledge is available about their actual ecological roles (Grossart et al. 2019). Saprotrophs participate in decomposition processes of dead organic matters, while parasitic fungi colonize living hosts to obtain nutrients for their life cycle (Donk & Bruning 1992). Parasitic fungi have a special significance because they can infect inedible phytoplankton species and facilitate energy transfer (during algal blooms in particular (Gleason et al. 2015)) to zooplankton via their zoospores (Kagami et al. 2007, Agha et al. 2016, Garvetto et al. 2019), a mechanism known as mycoloop (Kagami et al. 2014). In turn, parasitic chytrid outbreaks can be mitigated and suppressed by grazers who feed on their zoospores, channelling nutrients across trophic levels (Kagami et al. 2011, Frenken et al. 2020). However, the field of aquatic chytrid biology is highly skewed toward culture-based studies, especially ignoring DMF, which prevents the achievement of a more complete, detailed understanding of this fungal group (Laundon & Cunliffe 2021). In marine ecosystems, our understanding of fungal host–parasite systems is even more limited, potentially due to the challenges in isolating and culturing the fungal partner (Gladfelter et al. 2019), and in determining their ecological role that can vary under different circumstances (Grossart et al. 2015).

Aquatic fungi, and marine fungi in particular, have received much less attention in planktonic research compared to other planktonic microbes such as bacteria or protozoa (Amend et al. 2019). Despite that our knowledge on their diversity in marine ecosystems is greatly limited (Richards et al. 2015), a few studies have shown that distinct marine habitats harbour diverse and discrete fungal (mycoplankton) communities (Jeffries et al. 2016).

Specifically, coastal habitats constitute a transitional zone between riverine and open ocean (offshore) sites. These coastal habitats function as sinks for terrestrial-sourced fungi, and host higher proportions of the early-diverging fungal groups such as chytrids or Cryptomycota compared to oceans that are dominated by fungi belonging to Dikarya (Picard 2017, Hassett et al. 2019). Thus, coastal areas with high terrestrial influence should be hotspots for aquatic fungi. The high fungal diversity and the elevated proportion of chytrids in the Baltic Sea may support this hypothesis (Hassett et al. 2019, Rojas-Jimenez et al. 2019). Nevertheless, we have limited knowledge on how the transition from freshwater inlets towards offshore sites differentiate mycoplankton communities, and what processes (e.g., community assembly mechanisms) affect their abundances.

Metacommunity concept provides a framework to reveal important mechanisms that simultaneously shape (and maintain) biodiversity variation at regional scale (Leibold et al. 2004). These mechanisms include, for instance, selection by the environment (environmental filtering), dispersal-related processes (e.g., dispersal limitation, habitat connectivity) and stochasticity (ecological drift). To our knowledge, only a single study (Yang et al. 2021) aimed to estimate these processes in mycoplankton communities and found that their influences significantly differ along a river-sea transect, having more stochasticity in the coastal and offshore sites. In a recent study, Leibold et al. (2022) recognized the problem in the assumption that these community assembly processes act on all species and habitats similarly. Instead, they emphasized that sites and species can vary widely, thus, be influenced differently by the different assembly processes. Furthermore, the authors also emphasized the need to acknowledge the influence of biotic interactions on metacommunity patterns which was lacking in previous concepts. Joint species distribution models (JSDMs) have the advantage to provide such important insights into the drivers of variation in species distributions, and with this, estimate how the contributions of space, environment, and biotic interactions, driving metacommunity assembly, differ among sites and species (Ovaskainen et al. 2019, Poggiato et al. 2021, Leibold et al. 2022).

Positive (or negative) covariances between species’ distributions, however, could just as easily be explained by sharing (or not sharing) environmental habitat requirements as by the presence of true biotic interactions. To infer biotic interactions from such data, appropriate methods are needed to discriminate between biotic and abiotic environmental drivers of species distributions (Golding et al. 2015). To overcome this issue, probabilistic graphical model-based network approaches have been proven to successfully deal with indirect associations (as a result of shared environmental niches) by the incorporation of environmental metadata (Tackmann et al. 2019, Matchado et al. 2021). By revealing co-occurrences, networks have recently been useful to predict host–parasite interactions in virus- and chytrid-focused studies (Rojas-Jimenez et al. 2017, Kilias et al. 2020, Meng et al. 2021, Ilicic et al. 2022), even if network inference needs to be taken cautiously. Nevertheless, application of network approaches can help us to extend current knowledge on the relationships of fungi with other microorganisms (Rojas-Jimenez et al. 2017).

In this present study, we aimed to investigate fungal (mycoplankton) metacommunities of coastal habitats in a subarctic zone (Gulf of Bothnia, northern Baltic Sea, Sweden). The sparsely populated fungal reference databases and inconsistencies in the taxonomy provided by different reference databases can challenge the identification of fungal sequences from environmental samples (Nilsson et al. 2019). When different marker genes are targeted, this often leads to different classifications (Heeger et al. 2018). Therefore, in this study, we applied long-read metabarcoding approach using a Nanopore MinION sequencing platform which has been shown to provide accurate and efficient data in the characterization of aquatic environmental DNA (Davidov et al. 2020). Using recently developed novel joint species distribution model and network approach, we aimed to quantify the importance of biotic associations contributing to fungal metacommunity assembly, further, detect potential biotic (i.e., parasitic) interactions between fungi–algae pairs, respectively. We hypothesized a taxonomic transition of terrestrial to marine fungal species along the freshwater inlet–bay– offshore transects, with high fraction of early-diverging fungi, especially chytrids, in the bays. We also assumed that mycoplankton communities of bays should be influenced by a high degree of ecological drift and dispersal-related processes. Furthermore, co-occurrence network should reveal (i) chytrids that have strong co-occurrences with specific algal taxa, indicating putative, species-specific chytrid–algal parasitic interactions, and (ii) members of Cryptomycota as hyperparasites on parasitic chytrids.

## Materials and methods

### Sample collection

Samplings were conducted by following the course of four freshwater inlets towards the brackish water of the Gulf of Bothnia, northern Baltic Sea. Four surface water samples were collected (0.5 m depth) once in a month in these four bays during the summer period (June to September) in 2018 (Eriksson et al. 2022). In addition, samples from two offshore sites were also collected in each month (Fig. 1). All water samples were collected in sterile sampling bottles, then transported to the laboratory and stored in the dark at 4 °C. Physicochemical properties of samples were measured (Eriksson et al. 2022) and included in this study.

**Figure 1.**
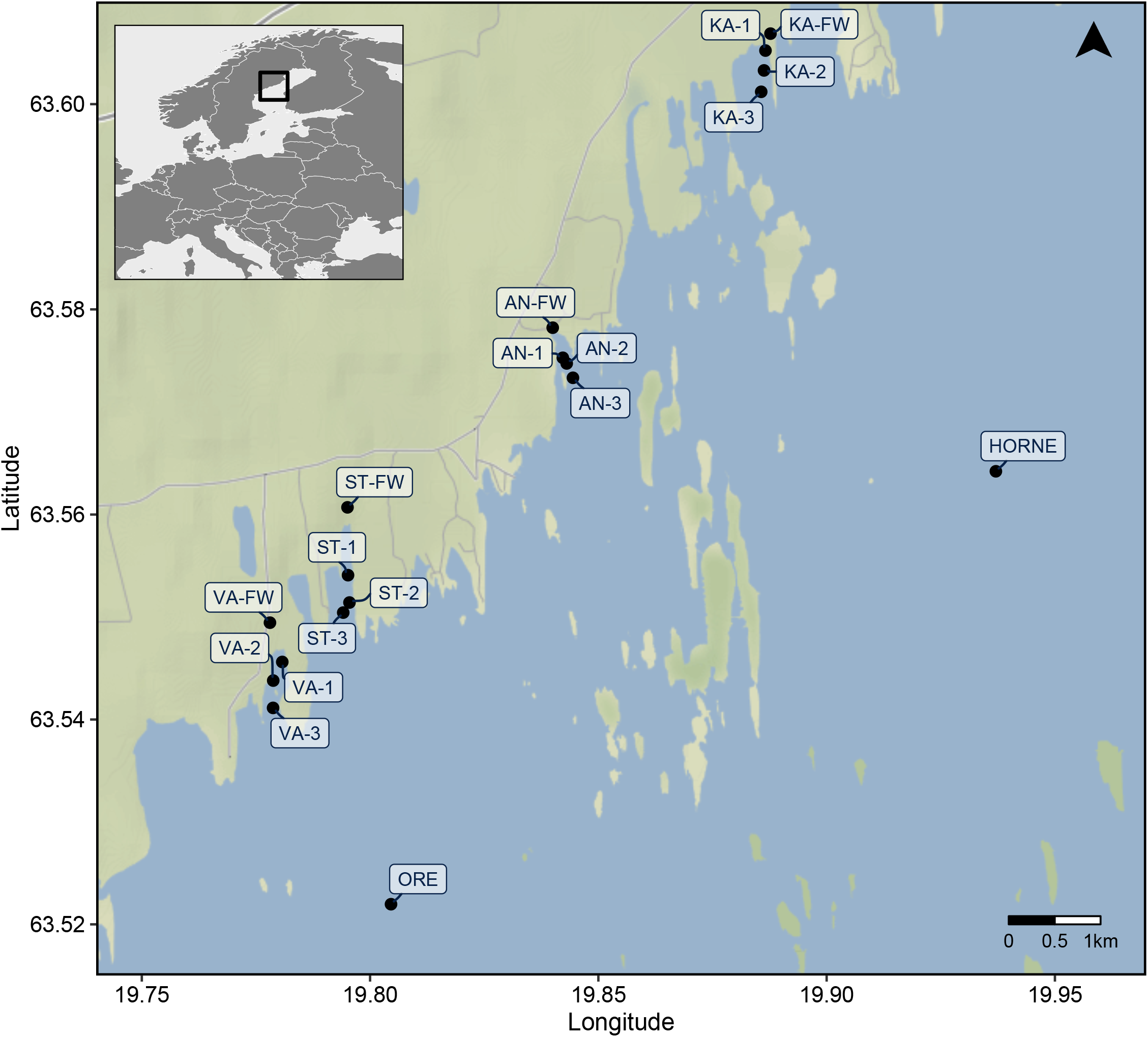
Four bays (KA: Kalvarsskatan East, AN: Ängerån, ST: Stadsviken, VA: Valviken) were sampled along the coastline of the Gulf of Bothnia, Sweden. FW refers to freshwater samples collected from the inlets, while numbers increase with distances from inlets. In addition, samples from two offshore sites (HORNE: Hörnefors, ORE: Örefjärden) were collected.

Briefly, temperature, pH and salinity were measured *in situ* with a WTW ProfiLine Cond 3110 portable device, and subsamples were used to measure environmental variables including dissolved organic carbon (DOC), total dissolved nutrients (TDP, TDN), dissolved inorganic nutrients (e.g., PO_4_^3-^, NH _4_^+^, NO_2_ ^-^, NO_3_ ^-^, SiO_2_), and humic substances.

Environmental data for the inlet (freshwater) samples of each bay is lacking. For the microbial community analysis, we aimed to filter 500 mL subsamples through sterilized 0.2 µm pore size membrane filters (Supor, 47mm, Pall Corporation) the same day as the water collection and kept the filters at –80 °C until further processing.

#### DNA extraction and sequencing

The DNA was extracted from the filters using the DNeasy PowerWater Kit (Qiagen) according to the kit protocol with some modifications. Namely, the samples were treated with an additional heating step (applying horizontal water bath for 30 min at 65 °C) and with a bead-beating step in each direction of 20 Hz, 3×3 min with a TissueLyser II (Qiagen) in order to aid the lysis of microorganisms including fungal cell walls. DNA extracts were quantified with a Qubit 4 fluorometer (Thermo Fisher Scientific).

To perform metabarcoding, we used the NS1short/RCA95m primer pair in order to amplify the majority of the ribosomal operon of the ribosomal tandem repeat (18S-ITS1-5.8S-ITS2-28S rDNA operon) (Wurzbacher et al. 2019). The PCR was performed, similarly as in Wurzbacher et al. (2019), but see details in Supplementary material. Amplification and its fragments length (∼4–6 kbp) were confirmed by gel electrophoresis resulting in 61 final samples (out of 72). The barcoded PCR products were purified with 0.8× of AMPure magnetic beads (Beckmann) following the manufacturer’s protocol. Thereafter, the purified PCR products were quantified using the Qubit 1× HS Assay Kit (ThermoFisher Scientific), pooled in equimolar amounts. This final pool was concentrated with 2.5× of AMPure magnetic beads in 50 µL nuclease-free water (ThermoFisher), and again quantified (Qubit 1× HS Assay Kit).

1 µg of library was used for the ONT library preparation using the 1D sequencing (SQK-LSK109; Oxford Nanopore Technologies). Briefly, amplicon library was end-repaired and adapted for nanopore sequencing using NEBNext Companion Module for Oxford Nanopore Technologies Ligation Sequencing (#E7180S), finally, a clean-up step was performed to enrich for amplicon length of > 3 kb using the Long Fragment Buffer (LFB) provided within the Ligation Kit. Sequencing was done on a MinION Mk1C instrument (ONT) operated with a Spot-ON Flow Cell (R9.4.1 chemistry). During the real-time monitoring, we noticed a low pore occupancy (e.g., high Strand:Single Pore ratio), hence, we prepared the remaining library with extended incubation times (+5 and +10 mins during end-prep. and adapter ligation, respectively) and loaded it to the existing run (55.14 fmol DNA in total). Real-time basecalling was executed using the High-accuracy basecalling (HAC) mode using the MinKNOW software (v21.05.12). In the end, 1.42 M reads were yielded with N50 read length of 4.4 kb and Q > 9.

#### Sequence data processing

Quality reads were demultiplexed and barcoded primers were trimmed with MiniBar (Krehenwinkel et al. 2019), and then filtered by length (3–7 kb) with NanoFilt (v2.8.0) (De Coster et al. 2018). These filtered, quality reads were then processed using the recently developed NGSpeciesID pipeline (v0.1.2.2) which wraps a set of tools to generate clusters and form polished consensus sequences for each cluster (Sahlin et al. 2021). It deploys the isONclust algorithm (Sahlin & Medvedev 2020) for read clustering (--mapped_threshold 0.8 --aligned_threshold 0.5) which accounts for variable error rates within reads. Draft consensus sequences were formed for each cluster containing at least five reads with spoa (v4.0.7) using maximum 500 sequences (--max_seqs_for_consensus 500), then reverse-complement clusters were merged using Parasail (v1.2.4). Finally, polishing of the consensus sequences (input reads are mapped back to the consensus sequence and basepair errors are corrected) were done with Racon (v1.4.20; Vaser et al. 2017) using two iteration steps. Since polished consensus sequences are the final output of the NGSpeciesID pipeline, we formatted the output files using custom scripts in R (R Development Core Team 2016) to have an appropriate input file for Mothur (v1.46.1; Schloss et al. 2009) to assign the corresponding sequence count to each OTU. We obtained on average 8,515 reads per sample with a mapping rate (percent of high-quality paired reads for generation of the consensus sequences from total reads) of 36.3 % (Supplementary Table S2). Sample coverage was assessed with the ‘iNext’ R package (Hsieh et al. 2016), and found that, with the exception of four samples, community composition was sufficiently covered (Fig. S1).

Taxonomic classification of the 512 final consensus sequences was done by local BLAST search against the nucleotide database (*nt*) from NCBI (downloaded on 23 October 2021) using BLAST+ (v2.11.0+), limiting the BLASTn search for Fungi (taxids: 4751) and keeping hits with at least 80 % identity. Results of the BLASTn search was processed with phyloR (https://github.com/cparsania/phyloR) to keep top hits (Supplementary Table S3) and to assign taxonomy levels. Unidentified (non-fungal) consensus sequences and their corresponding OTUs from the OTU table were filtered out. In the end, 504 OTUs were identified as fungi (with an average alignment length of 2,107 bp and 93.2 % identity), thus, were kept for downstream analyses. OTUs without matches below Kingdom level were classified as ‘likely fungi’. Quality-filtered (Q > 9) reads were deposited to NCBI SRA database under the accession number PRJNA849821.

#### Data analyses

Venn diagram was used to visualize the number of shared fungal taxa across sampling sites, furthermore, diversity analyses (alpha-diversity and beta-diversity based on Bray-Curtis distance) were performed using the ‘microeco’ R package (v.0.6.5) (Liu et al. 2021) and the results (i.e., nonmetric multidimensional scaling – NMDS) were plotted using ‘ggplot’ package (Wickham 2009). Difference in alpha diversity across bays, their inlets and offshore sites were tested with ANOVA followed by Duncan’s test (p < 0.05) as a post-hoc test. To test compositional differences between samples, permutational multivariate analysis of variance (PERMANOVA) with 999 permutations was performed using the function *adonis2* in ‘vegan’ R package (Oksanen et al. 2016). Distance-based redundancy analysis (dbRDA) was also performed on the bay samples to assess the influence of environmental variables or sampling time (i.e., Day). To statistically test their influences, Mantel test (Spearman correlation with 999 permutation) were computed on community dissimilarity based on Bray-Curtis distance.

#### Inference of the internal structure of metacommunities using joint species distribution modelling

To predict metacommunity structure as a whole, a recently developed scalable joint species distribution model (sjSDM) approach (Pichler & Hartig 2021) was applied which use Monte Carlo integration of the JSDM likelihood together with elastic net regularization on all model components. Due to the lack of measured environmental data for freshwater inlets and offshore sites, we used only samples from the four bays (n = 40, no. of OTUs = 451). Spatial eigenvectors were generated from the GPS coordinates to account for spatial autocorrelation and measured environmental variables were z-transformed prior the analysis. The regularization for all covariances and coefficients (following general suggestions from the developers) was tuned over 40 random steps with leave-one-out cross validation (LOOCV), 150 iterations and learning rate of 0.01 using the *sjSDM_cv* function of the sjSDM R package (v1.0.1) (Pichler & Hartig 2021). Thereafter, the best regularization parameters were used to fit a multivariate probit model using the *sjSDM*.*tune* function. The model is available as Supplementary R Data file.

Following the framework by Leibold et al. (2022), our model was then used to estimate how the contributions of environment, space and biotic associations shaping metacommunity assembly vary among sites and taxa. This quantitative approach allowed us to identify how different sites and taxa contribute to the overall metacommunity structure, taking into account the fact that a set of sites or taxa are not necessarily equally influenced by the interplay of environmental filtering (abiotic selection), dispersal, biotic interactions and ecological drift (Leibold et al. 2022).

#### Network analysis

With the aim to further investigate biotic interactions and, in particular, potential parasitic-host associations, co-occurrence network was constructed. In parallel to this study, 18S V6-V8 rRNA gene sequences have been generated using the same samples (Eriksson et al. 2022) and the subset of that dataset was used to assess potential host–parasitic(saprotrophic) associations in our bay samples (n = 40). Absolute read counts were used for the network analysis because previous studies have shown that relative abundance data suffer from apparent correlations which lower specificity of association networks (Berry & Widder 2014, Meng et al. 2021). Further, since low number of sites are susceptible to false positive correlations (Berry & Widder 2014), we decided to run one global network analysis without subsetting our dataset into the three time periods or to individual bays. Nevertheless, in this analysis, we used only the most dominant fungal groups (Cryptomycota and chytrids) that are prone to have parasitic lifestyle, but also included the ‘likely fungi’ group to assess their putative cross-kingdom associations. To account for variations in sequencing depth between the long-read fungal and 18S-based algal datasets, we decided to use FlashWeave (v0.18.1) (Tackmann et al. 2019) which applies *clr* transformation to handle compositionality with its adaptive pseudo-counts (*clr-adapt*) approach. Prediction of ecological interactions (alpha < 0.05) between fungi and algae was performed using FlashWeave-S (‘sensitive’ mode) with default settings. For enhanced reliability, associations were computed only when an OTU was present more than 20 times (automatically determined by the software). The integration of 15 meta-variables (MVs such as ‘Day’ as sampling time, ‘Bay’ as identities, and measured physicochemical variables) was done in order to remove potential indirect associations (i.e., as a result of shared niche preference). The constructed cross-kingdom network was visualized with Gephi (v0.9.2) using associations’ weight cutoff > 0.3 and the OpenOrd layout. For clarity MV nodes were moved to the side.

All statistical analyses were performed in R v4.0.4 implemented in RStudio v1.4.1106. For the analysis of JSDM, the Python package ‘PyTorch’ (Paszke et al. 2019) was run from within R thanks to the ‘reticulate’ R package (Allaire et al. 2018). Julia (v1.6.4) was used for constructing association network with FlashWeave package (Tackmann et al. 2019) from within the ‘microeco’ R package (Liu et al. 2021).

## Results

### Fungal diversity and community structure

Our long-read (18S-ITS1-5.8S-ITS2-28S rDNA) metabarcoding pipeline revealed 504 fungal OTUs. Freshwater inlets of Valviken (VA; n = 9), Ängerån (AN; n = 8) and Kalvarsskatan East (KA; n = 4) harbored most of the unique taxa (OTUs), while the offshore sites, Hörnefors (HORNE) and Örefjärden (ORE), did not have any site-specific fungus (Fig. S2). In total, 113 fungal taxa were shared between all sampled sites. Generally, the alpha diversity (based on Shannon and Simpson indices) declined significantly (*p* < 0.05) from the inlets towards the offshore sites (Fig. 2).

**Figure 2.**
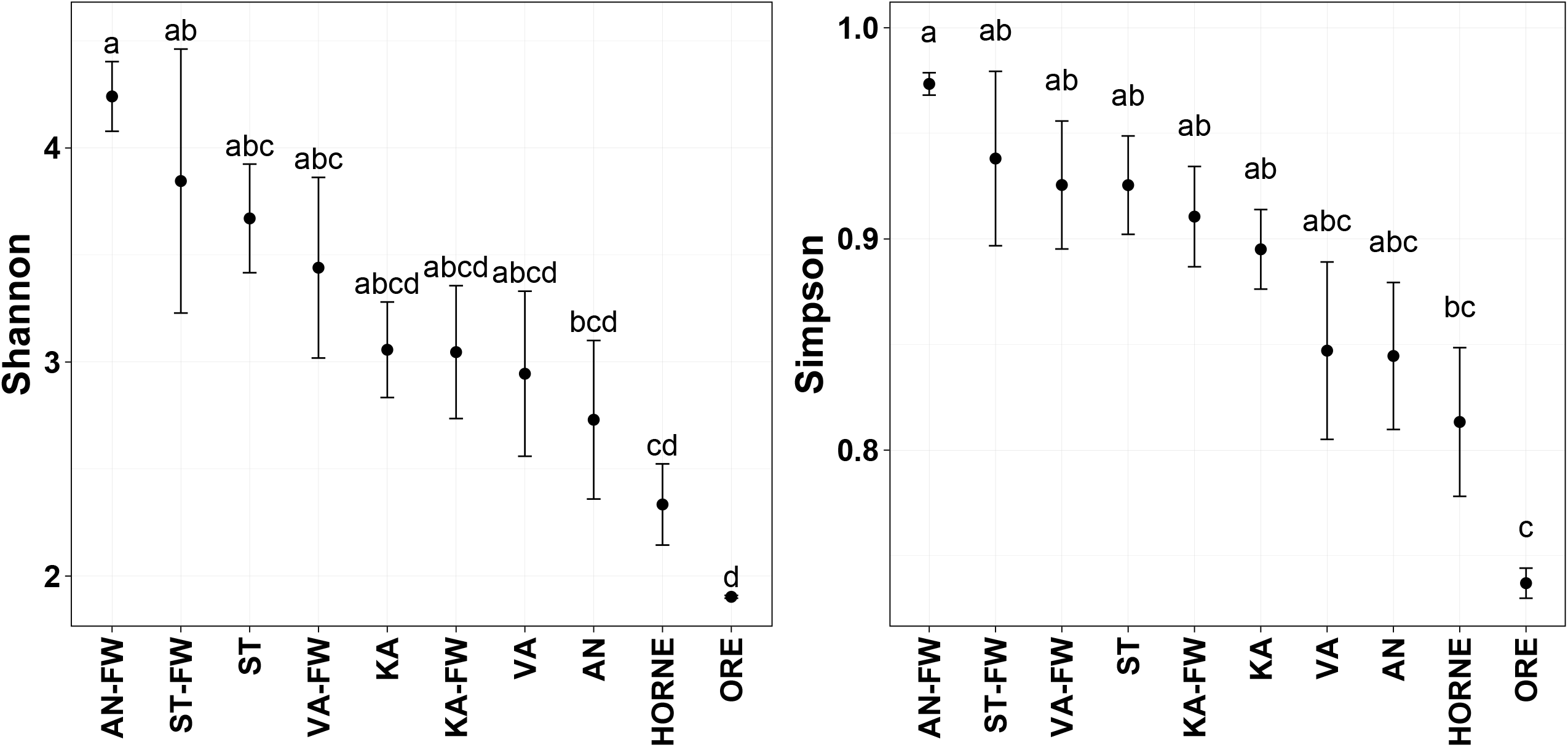
Alpha-diversity measures (Shannon and Simpson) for the four bays, their respective freshwater inlets and the offshore sites. For IDs reference, see Figure 1. Significant post-hoc groups (*p* < 0.05) among the different sites are represented by lowercase letters.

Early-diverging fungal groups such as Cryptomycota and Chytridiomycota represented the most dominant fungal groups in our samples (Fig. 3), with 93 and 133 OTUs, respectively. Their relative distribution, however, varied greatly across sampling sites. For instance, the relative abundance of Cryptomycota was highest (15.3–68.7 %) in freshwater inlets and lowest (3.5–6.9 %) in offshore samples, moreover, decreased along the north-south gradient (KA → AN → ST → VA). In contrast, Chytridiomycota fungi increased their relative abundance towards bays (18.7–52.0 %) and offshore sites (37.6–54.8 %) with the exception of ST (= Stadsviken; 22.5 %) and its inlet (ST-FW; 31.6 %). Relative abundance of fungi assigned as Ascomycota decreased from inlets (14.1–26.4 %) towards offshore sites (0.9–1.6 %). Basidiomycota, on average, represented only 2.8 % of the fungal communities and they were more common in bays and their inlets (3.2 % and 3.75 %, resp.) than in the offshore sites (0.3 %). Unassigned fungi (‘likely fungi’) accounted a great portion in almost all communities (26.9 % in average) and their portion increased in the offshore sites up to 53.6 %. Other fungal groups such as Microsporidia (0.3 %), Mucoromycota (0.05 %), Zoopagomycota (0.03 %), Sanchytriomycota (< 0.01 %) and Blastocladiomycota (< 0.01 %) can be considered as rare fungi in these aquatic habitats with neglectable mean relative abundances.

**Figure 3.**
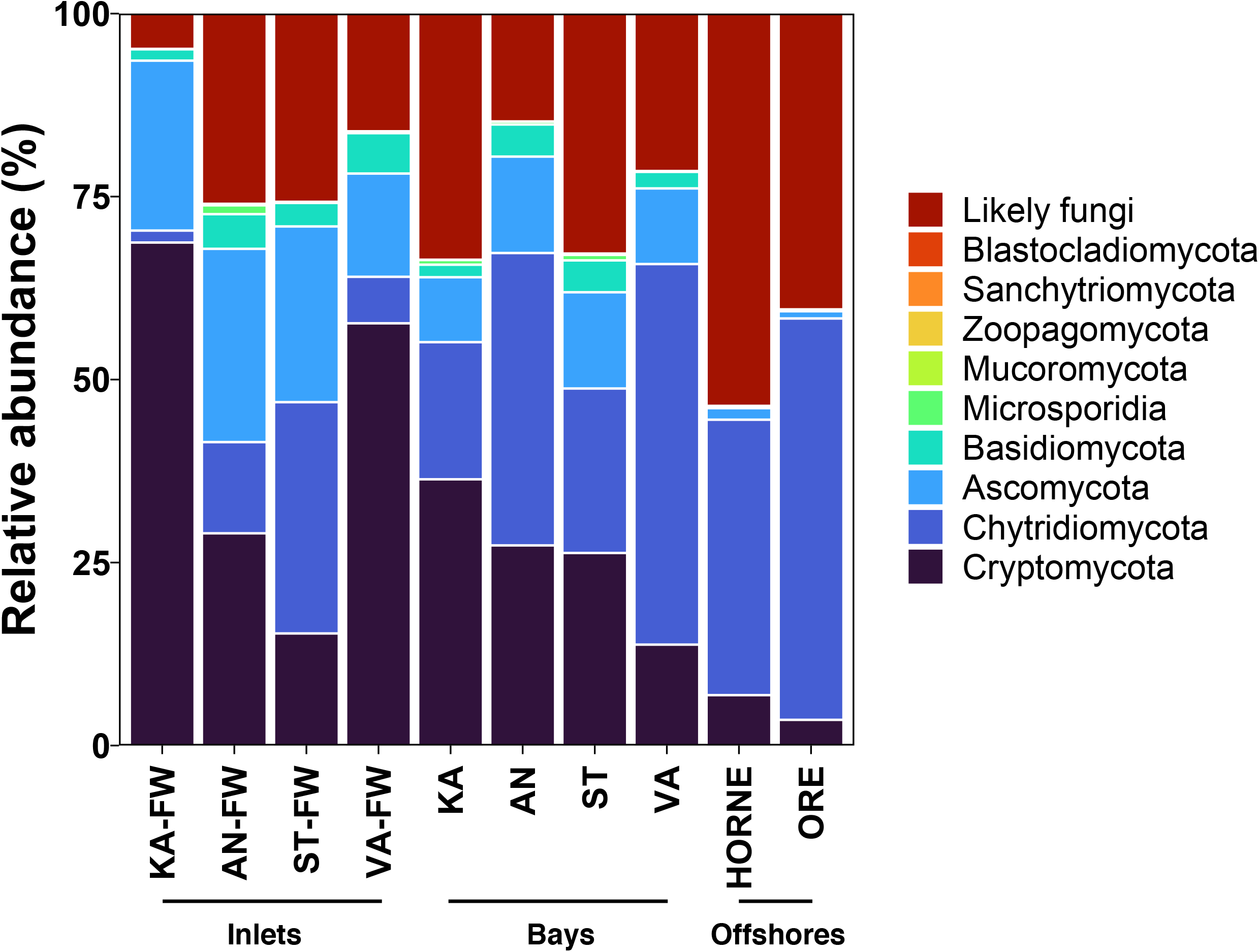
Relative abundance of fungal phyla across the bays, their freshwater inlets and the two offshore sites. ‘Likely fungi’ refers to OTUs that were assigned as fungi (based on BLAST search) but without matches to any phyla.

Fungal community compositions exhibited strong compositional shift over time (from June to September, F = 9.90, *p* = 0.001), although the PERMANOVA test also confirmed spatial variability (F = 2.28, *p* = 0.001) which was mainly pronounced by the separation of bay samples from their respective freshwater inlets (Fig. 4a). The significant interaction between sampling sites and time (F = 1.95, *p* = 0.001) further showed that temporal variation differed between sites. When the fungal metacommunities of the four bays were analysed alone (using dbRDA) to investigate the influence of the measured environmental variables (Fig. 4b), temperature, pH, humic substances, salinity, DOC, NH_4_^+^, TDN, and, to a lesser degree, NO_3_ ^-^ and chlorophyll-*a* concentrations were important (all *p* < 0.05) in shaping the beta diversity of fungal metacommunities, besides sampling time (i.e., Day) (see the results of Mantel test in Table S4). The three distinct clusters are clearly apparent from the dbRDA (Fig. 4b) and accord with sampling time rather than with space (i.e., bays) when environmental parameters were considered.

**Figure 4.**
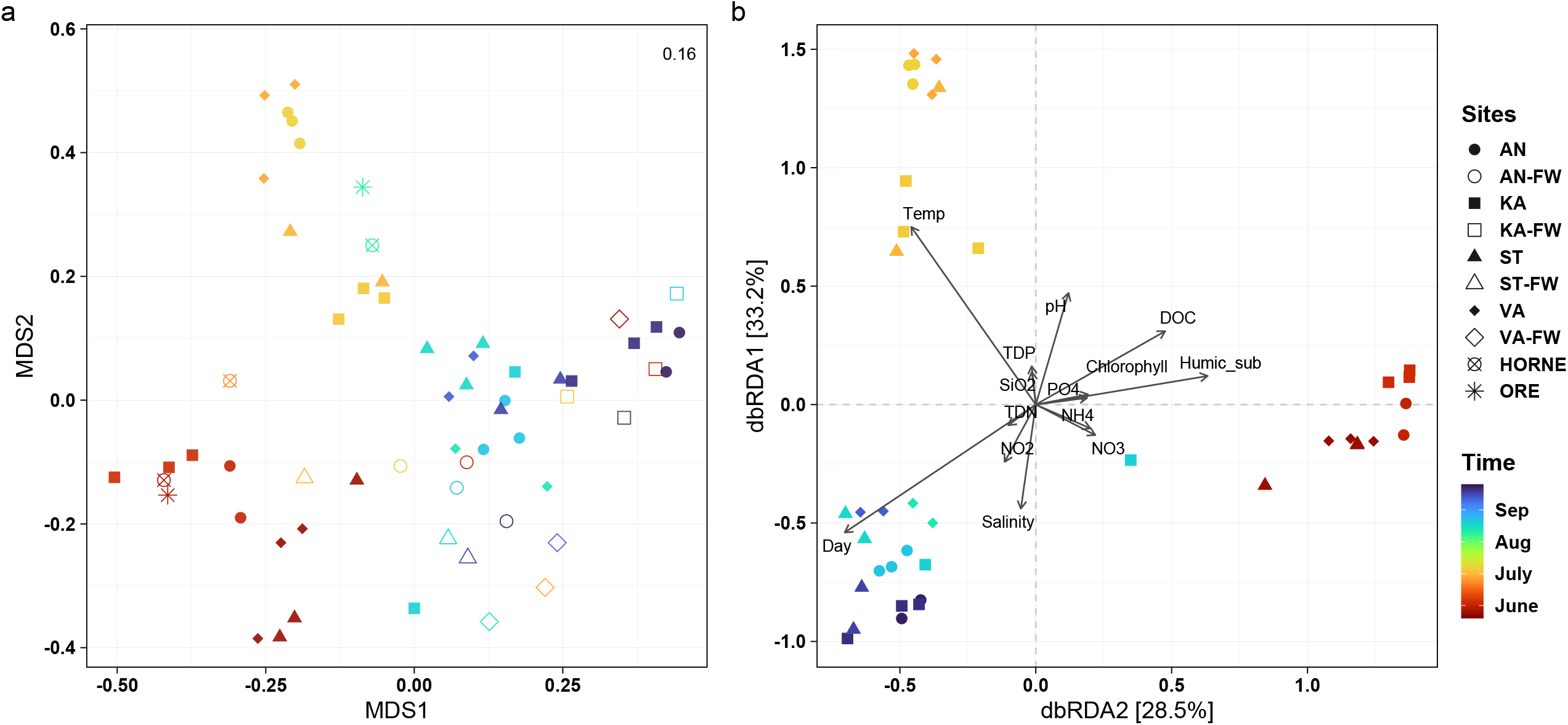
(a) NMDS plot (based on Bray-Curtis distance) shows differences in fungal community structures across all sampling sites (four bays, their inlets and two offshore sites), coloured by sampling time (June–September). Stress value is shown on the upper right corner. (b) Distance-base redundancy analysis (dbRDA) plot reveals the influence of environmental conditions across bays and the distinct clustering by sampling time.

These temporal dynamics in fungal metacommunities can be attributed to several trends observed in the relative abundances of different taxonomic groups (see Fig. S3). Specifically, Cryptomycota (e.g., *Paramicrosporidium* sp. and mainly unidentified Cryptomycota (73.1 %)), Ascomycota (members in the genera of *Articulospora, Cladosporium, Penicillium*), Basidiomycota (*Vishniacozyma* sp., *Rhodotorula* sp.) and Microsporidia (*Mitosporidium* sp.) showed elevated abundances towards late summer (August and September), while Chytridiomycota had its peak in July (dominated by OTUs assigned as *Betamyces* spp. and *Chytridium polysiphoniae*) and decreased similarly as ‘likely fungi’ by the end of the sampling campaign (September). In the remaining fungal groups, there was no clear, detectable trend.

#### Internal structure of aquatic fungal metacommunities

We used scalable joint species distribution model to estimate the importance of environment, space and biotic interactions in driving metacommunity assembly among sites and fungal taxa of the bays. Our model revealed that the distribution of fungal taxa was mainly driven by the joint effects of space and environment (Fig. 5, Table S5). Sites within each bay differed greatly in how their community compositions were attributed to environmental and spatial effects, as well as biotic associations (Fig. 5, left panel; Fig. S4), but were explained by similar average R^2^ values (approx. 1.2 %). Community-level variation in each bay showed enhanced importance of biotic interactions (or co-distributions) in particular of sites which were occupied by taxa with non-extreme optima. This pattern occurred mainly in samples collected in the months of July and September (except in the case of KA whereas June samples also showed enhanced effect of biotic component) (Table S5).

**Figure 5.**
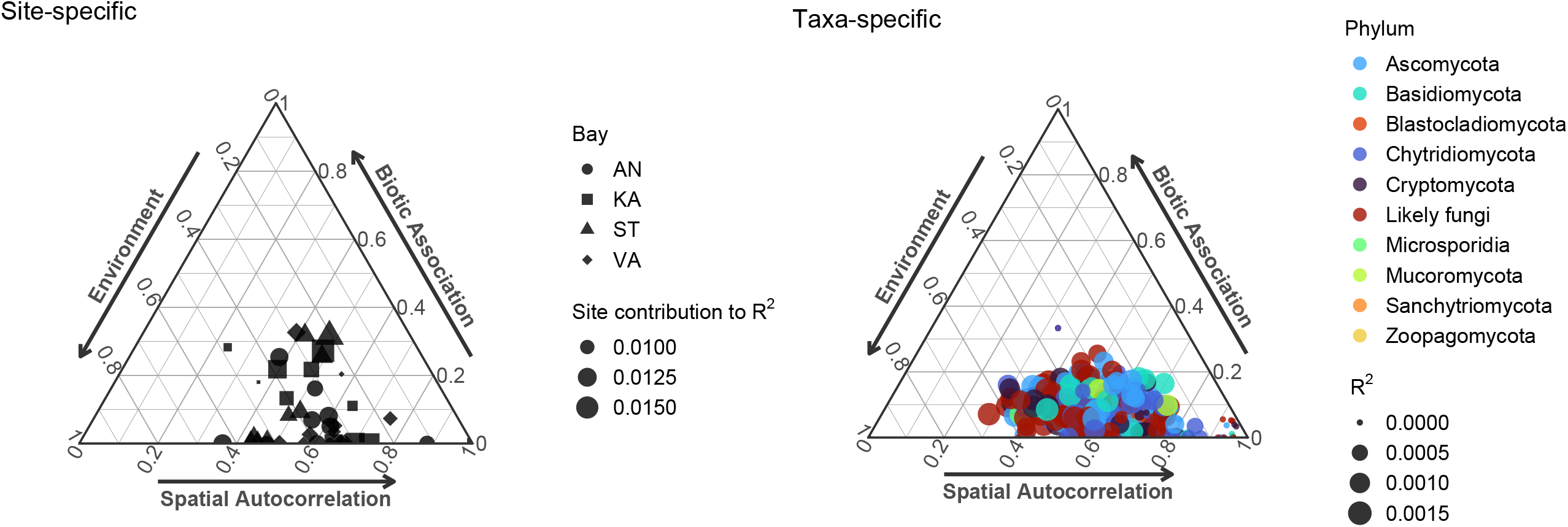
Site-specific (left panel) and taxa-specific (right panel) internal structure of fungal metacommunities of the bays assessed by a scalable joint species distribution model (sjSDM). The relative influence of environment (e.g., through environmental filtering), space (e.g., dispersal limitation) and biotic associations was estimated on the distribution of mycoplankton metacommunities. The size of the symbols indicates the amount of variation explained (R^2^) by the model for each OTU (n = 451) or site (n = 40).

Taxa distributions were weakly predicted by the model (e.g., low R^2^ values) and were slightly greater influenced by biotic associations as the effect of the environment and space decreased (Fig. 5, right panel). We also found that the model resulted in even lower R^2^ values when taxa were almost exclusively influenced by space and covered broad niches (i.e., when environmental filtering was weak). OTUs within each dominant phylum were distinctly affected by the three components (Fig. S5), hence, the distribution of each taxon was determined by a unique combination of environment, space and biotic factors. Phyla that are represented by a very few members (i.e., Blastocladiomycota, Mucoromycota, Sanchytriomycota and Zoopagomycota) and considered as rare were influenced by spatial factors to a great extent, while Microsporidia occurrences were mainly affected by the environmental conditions. It is, however, important to mention that temporal factor (i.e., Day), despite its strong influence as presented in the above-mentioned multivariate analyses, was not included in the model in order to investigate the pure effects of measured environmental variables on taxa distributions.

#### Co-occurrence network

We generated one cross-kingdom co-occurrence network on all bay samples to predict ecological interactions between specific fungal groups (Cryptomycota, chytrids and ‘likely fungi’) and algae (Fig. 6, S6). In total, this final network was composed of 303 nodes (288 OTUs) and 498 associations (edges). Detailed list of all interactions can be found in Supplementary Table S6. Among the three fungal groups (Cryptomycota, Chytridiomycota and ‘likely fungi’) 152 interactions were detected, of which 90.7 % were positive. The strongest associations (e.g., corr. weight > 0.9) occurred between OTUs within the same taxonomic group. Between Cryptomycota and Chytridiomycota eight associations (two negative and six positive) were found, and these chytrid OTUs had no link to any algal OTUs. To reveal potential parasitism (corr. weight > +0.3) between kingdoms, we found 40 interactions. Most (19) were identified between chytrids and algae (eight Ochrophyta, five Chlorophyta, four Cryptophyta and only two Dinoflagellata). The five strongest interactions showed links to: a Dinoflagellata (Syndiniales) (corr. weight = 0.71), three Chlorophyta identified as *Chlorella* sp. (0.66), *Chlamydomonas* sp. (0.63), *Cladophora glomerata* (0.53), and an OTU belonging to Dinophyceae class (0.61). Interestingly, the group of ‘likely fungi’ had 12 associations with algae with similar taxonomic distributions (five-five Ochrophyta and Chlorophyta, but only one-one Cryptophyta and Dinoflagellata). Within this group, the strongest interactions here occurred with *Koliella* sp. (0.55), *Chaetoceros* sp. (0.44) and an unassigned Chrysophyceae (0.40). Cryptomycota had the least number of positive edges with algal OTUs (nine) and those algae mostly belonged to Dinoflagellata (e.g., *Dinophysis* sp., *Prorocentrum* sp. and *Levanderina* sp.).

**Figure 6.**
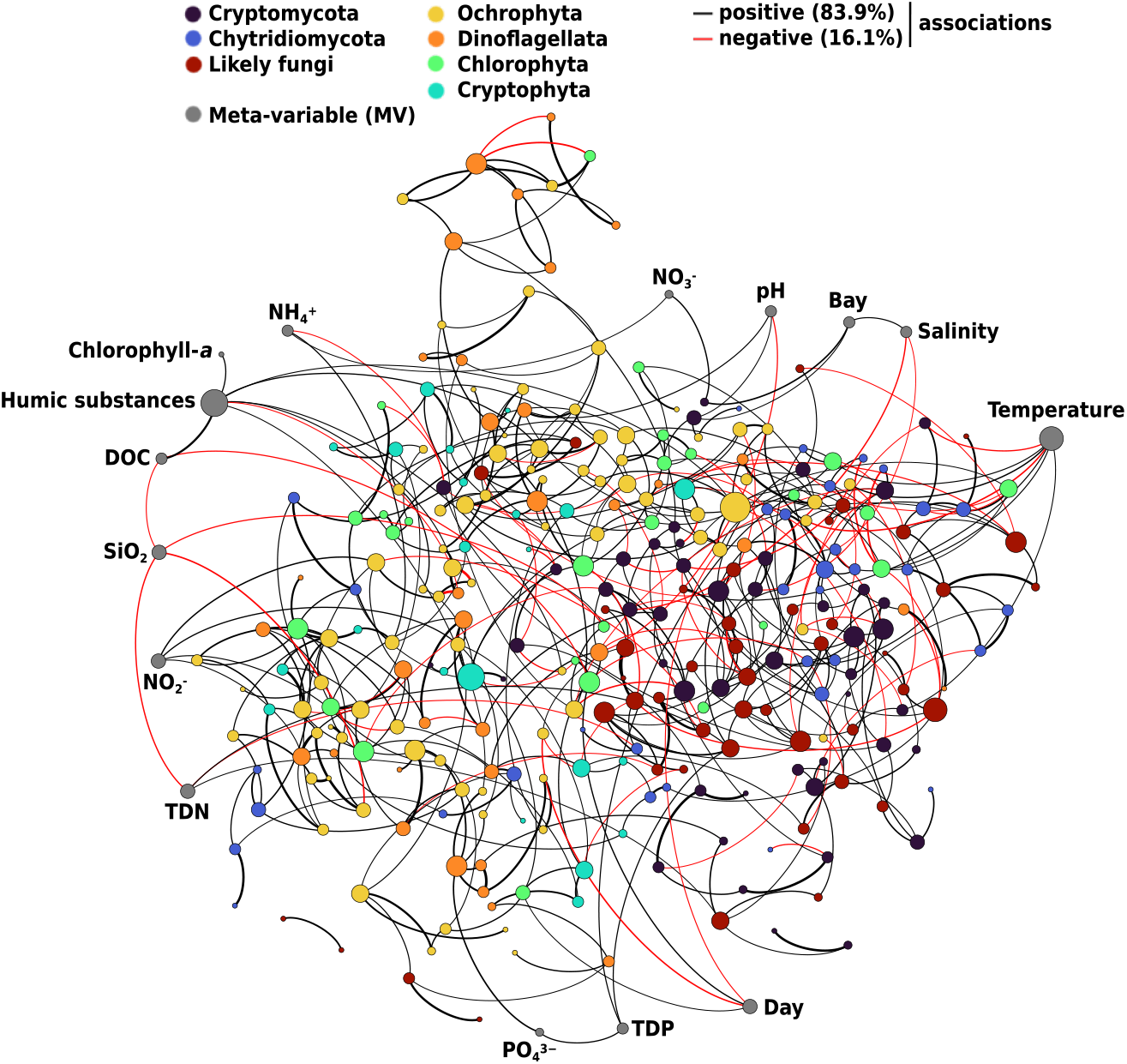
Prediction of fungi–algae interactions with FlashWeave (alpha < 0.05). The size of nodes (OTUs or meta-variables; n = 303) are proportional to the number of predicted interactions (degree). Edge width refers to the strength of correlations (weight cutoff > |0.3|; n = 498) and coloured by association type (positive or negative). ‘Likely fungi’ refers to OTUs that were assigned as fungi (based on BLAST search) but without matches to any phylum.

MVs, such as measured environmental parameters and/or sampling time can lead to spurious correlations between OTUs that share similar niche optima and thus associated with the same MV. The inclusion of MVs in our network analyses showed that several fungal (15) and algal (25) OTUs directly associated to the included MVs. The most influential were water temperature (with seven links) and humic substances (with six links). The former influenced four fungal OTUs and three algal taxa, while the latter had only links with algae (mainly Ochrophyta). Sampling time (Day), interestingly, only associated with four algal taxa. More precisely, *Nannochloris* sp. and *Pyramimonas* sp., both within the group of Chlorophyta, showed positive associations (e.g., positive trend in their abundances over the sampling course), while *Diatoma* sp. and *Uroglena* sp. had both negative associations. There were two Cryptomycota OTUs showing positive association with sampling sites (Bay) and these OTUs were also highly influenced (71–75 %) by spatial effects as revealed by sjSDM.

## Discussion

Our long-read metabarcoding study identified 504 fungal taxa along four transects encompassing freshwater inlets up to their corresponding bays and beyond to offshore habitats in the Gulf of Bothnia, Sweden. A great fraction of the mycoplankton (44.8 %) was assigned as early-diverging fungi (i.e., Cryptomycota and Chytridiomycota) which aligns with previous studies that reported the dominance of these fungal phyla in both freshwater and marine environments (Comeau et al. 2016, Hassett & Gradinger 2016). Generally, alpha diversity of mycoplankton declined and community compositions changed along these four transects (from freshwater to offshore sites). This resonates with a general declining trend found by Yang et al. (2021) in a much greater spatial scale along the Elbe River down to its estuary (North Sea). As we hypothesized, chytrids showed elevated richness in bays and offshore sites, and abundances in mid-summer (i.e., July). This is the first study, to our knowledge, that discusses the relevance of spatiotemporal shifts in the composition of coastal mycoplankton metacommunities, moreover, highlights the varying contributions of both sites and taxa to their metacommunity structures. Our network analysis showed a high number of co-occurrences among the dominant fungal groups (Cryptomycota, Chytridiomycota and ‘likely fungi’ that includes all unassigned fungal-like OTUs), suggesting a great proportion of biotic interactions. Further, the 40 strong associations in fungal–algal OTUs indicates putative host–parasite/saprotroph links that can provide basis for further investigations.

### Composition and diversity of coastal mycoplankton

Our study shows that early-diverging fungal phyla dominate not only sheltered aquatic ecosystems, as previously suggested (see e.g., Rojas-Jimenez et al. 2017), but also in habitats which are greatly affected by terrestrial input and continuous mixing events. Dominance of chytrid-like sequences was also found in a field survey, covering a great selection of marine (Arctic) and freshwater (temperate biomes) environments (Comeau et al. 2016). Their results, however, showed much less frequency and abundance of Cryptomycota, in contrast to our findings. The general compositional shift over sampling sites differed from the one found in the study of Yang et al. (2021). Namely, the authors found greater dominance of chytrids in samples with greater freshwater influence and the dominance of Ascomycota and Basidiomycota in open sea (offshore) environments, while our results suggest the opposite.

This disparity may arise due to a greater salinity range their study covered, increased anthropogenic activities around that study area (i.e., influence of the city of Hamburg), or due to the lack of the inclusion of temporal aspect in their study. All these might affect mycoplankton communities in space, creating distinct community structures and disparities among studies. Picard (2017) has found greater dominance of Ascomycota in mycoplankton communities in coastal North Carolina, and chytrids were more dominant fraction of the fungal communities in sediments, suggesting the importance of dormancy (Velasco-González et al. 2020). Previous studies targeting aquatic microfungi are scarce, but the findings so far indicate that the proportion of different fungal groups can greatly dependent on spatial scale and the environment they inhabit.

In spite of the high diversity of ‘dark matter fungi’ in aquatic environments (Grossart et al. 2015), we lack detailed taxonomic information for most Cryptomycota OTUs (73.1 %) in this present study, although their ecological importance is most likely relevant. This may not be surprising, giving the fact that newer clades and taxa are continuously discovered and described, thus, the taxonomy of such early-diverging fungal lineage is not completely resolved. Among Cryptomycota, an intranuclear parasite of amoebae (*Paramicrosporidium saccamoebae*) occurred in most freshwater inlets, further, with an elevated abundance during August and September in the bays. This indicates that even though freshwater inlets could nearly continuously support dispersal of parasitic Cryptomycota into coastal and offshore habitats, their establishment is most likely determined by other factors (e.g., hosts presences). Interestingly, two occurring *Rozella* spp. (commonly described as hyperparasites) did not end up in the co-occurrence network, suggesting the lack of hyper-parasitism and/or algae parasitism of these taxa in the observed bays.

*Betamyces* spp. and *Chytridium polysiphoniae* were dominant members of chytrids in July and in the bays, particularly, and have been reported as parasites on several phytoplankton species (Christaki et al. 2017) and on seaweed in estuarine and marine habitats (Gleason et al. 2011). This, in line with our hypothesis, further supports that coastal habitats provide an optimal environment for chytrids (Hassett et al. 2019) whereas the mixing of riverine and open ocean samples promotes the growth of phytoplankton and source aquatic fungi with parasitic lifestyle. The peak abundances of chytrids in July indicates the presence of parasitic taxa which usually emerge in the presence of elevated algae biomass, channelling nutrients to higher trophic levels via the mycoloop (Grami et al. 2011, Gerphagnon et al. 2015, Frenken et al. 2017).

Phylum Ascomycota was dominated by a member of the genus *Articulospora* (namely, *Articulospora atra*) which is a commonly known aquatic hyphomycetes, identified in numerous freshwater ecosystems, previously. To our knowledge, this is the first record showing that it can establish great population size (as a proxy of its relative abundance) in brackish ecosystem, likely dispersed from the freshwater inlets into the bays. *Nectria cinnabarina* (also among the top 50 taxa) is a weakly parasitic and then saprotroph fungus on deciduous hardwoods. Within this genus some anamorphs such as *Heliscus lugdunensis* and *Flagellospora curta* are, for a long time, known as aquatic hyphomycetes (Webster 1959, 1992). Since the link between anamorph-teleomorph life-history stages are still poorly known (i.e., only 10 % of described taxa) (Bärlocher & Marvanová 2010), we cannot rule out the possibility that our detected *Nectria* taxon might be just another fungus with the capability to growth asexually in aquatic habitats. Numerous members of Ascomycota and Basidiomycota (e.g., *Cladosporium* and *Blumeria*) found in this study are known as fungi with terrestrial origin. These taxa potentially represent nutrient sources for saprotrophic aquatic fungi (Magyar et al. 2016). However, more and more evidences suggest that a fraction of these fungi display a truly amphibious ability, hence, their presence as mere metabolically inactive spores or relictual DNA is questioned (Amend et al. 2019). A recent study suggests the role of terrestrial (i.e., phylloplane) fungi in the decomposition of submerged litter (Koivusaari et al. 2019). Their results pointed out that the majority of their observed fungi in streams had both plant- and water-associated lifestyle, moreover, they have found fungi (within Basidiomycota) which were active but rarely observed in aquatic (lotic) system. Similarly, Rojas-Jimenez et al. (2019) speculated that fungi belonging to Ascomycota, Basidiomycota and Zygomycota may participate in decomposition processes, or even act as biotrophs of phytoplankton, other microbes (i.e., protozoa) and microinvertebrates. Taken together, future studies should go beyond aquatic fungi and investigate systematically the ecology and ecosystem functions of other fungal groups with known terrestrial origin.

The low taxonomic recovery of aquatic fungi (especially zoosporic fungi) also suggests the limitation of even the biggest reference databases that likely cannot assign adequately (semi)-aquatic taxa to taxonomic levels below phylum-level. This limitation further enhanced when we found no overlap between our community compositions and a recent study that aimed to reveal the diversity of marine fungi in the Baltic Sea (Tibell et al. 2020). Although, they identified species mostly by morphology. The closest match (i.e., *Leptosphaeria biglobosa*) occurred with a set of *Leptosphaeria* species which, decomposing wood substrates, apparently a diverse group (besides *Corollaspora* spp.) in the brackish environment of the Baltic Sea. Nevertheless, similar issues must be resolved in the near future by complementing culture-based studies with metabarcoding approaches in order to improve the currently sparsely populated fungal databases (Nilsson et al. 2019).

### Metacommunity dynamics

Besides the major impact of salinity in shaping fungal communities (Rojas-Jimenez et al. 2019, Ilicic et al. 2022), our results highlight the relevance of other abiotic factors (i.e., temperature, pH, humic substances, etc.) as well as the influence of time in affecting community dynamic of mycoplankton. This emphasizes that snapshot studies easily can miss important aspects of community dynamics, which are clearly influenced by the simultaneous influence of space and time. In general, the importance of assembly processes shaping aquatic fungal communities is largely unknown. To our knowledge, only Yang et al. (2021) aimed to estimate community assembly processes and found that selection was more important in less saline environments (freshwater inlets) together with dispersal limitation, while communities of the coastal and offshore sites were mainly assembled by mass effect and ecological drift. Modelling the internal structure of metacommunity helped us predicting the occurrences of each mycoplankton member and estimating the relative importance of abiotic and biotic factors that affect their distributions in the observed bays. Our results suggest that the observed bay-inhabiting microfungi had occupied rather broad niches (based on our measured environmental variables) but were potentially limited by dispersal as evidenced by slightly greater importance of spatial effects than environment. For example, a parasitic chytrid, *Quaeritorhiza haematococci*, was unique to the bay of Kalvarsskatan East and an uncultured Cryptomycta in the bay of Valviken. Rojas-Jimenez et al. (2017) showed, similar to this present study, the influence of spatial factors (“site-specific effects”) in fungal communities of ice-covered lakes, indicating that the detected mycoplankton composition were highly dependent on the origin of samples, both across the landscape and within a single habitat. In extreme cases when taxa were highly influenced by space, their distributions were predicted very weakly by the model as the low R^2^ values suggest. As our findings show, taxa-level distributions of mycoplankton are generally much more difficult to predict compared to community-level predictions, potentially due to other factors (i.e., trophic interactions, unmeasured environmental parameters) that were not included in our model. Therefore, these results must be considered cautiously. This may align with Leibold et al.’s (2022) simulation results that suggested greater predictive power for species with narrower (more specialized) environmental niche than those that are dispersal limited, thus, show high spatial effects. Our overall findings suggest that both environmental and spatial factors can easily enhance processes (i.e., selection driven by abiotic environmental factors and/or dispersal-related mechanisms) that contribute to the internal structure of metacommunities, and eventually determine the distribution of microfungi. Nevertheless, the inclusion of biotic associations further highlighted that their importance can strengthen when the abiotic and spatial drivers are not dominating. It is important to note that the biotic component of the model (i.e., biotic associations) might mask processes resulting from ecological drift (stochasticity), while spatial patterning might include unmeasured but spatially autocorrelated environmental variables (Blanchet et al. 2020, Wilkinson et al. 2021). In field studies, therefore, the presence of complex trophic interactions (i.e., viruses, grazers), unmeasured environmental variables may all represent additional challenges in teasing apart processes that structure microbial communities, and thus, cause the inappropriate detection of the effect of environment–space–biotic components in this particular joint species distribution model applied here. Although our results must be discussed with care, we believe that, by taking into account interactions (or more precisely covariances) between species, joint species distribution models can represent a useful tool for researchers who aim to investigate distributional patterns.

### Putative biotic interactions

Our network analysis revealed a diverse picture, with a high number of co-occurrences (138) among three fungal groups (Cryptomycota, Chytridiomycota and ‘likely fungi’) which support the results found by sjSDM that estimated in average 5.48–7.13 % contributions of biotic associations among these groups. This highlights that the co-occurrences identified by network approaches can indicate direct pairwise interactions, although it might also suggest co-distributions via a complex interplay between true biotic associations, unmeasured environmental factors and, naturally, ecological drift (stochasticity).

Numerous associations (40) between fungi and algae were also discovered. Interestingly, one of the most abundant chytrids (*Betamyces* spp.), which are recognized as parasites on numerous phytoplankton species (Christaki et al. 2017), showed strong associations with only an unassigned Dinophyceae taxon and a chrysophyte (*Paraphysomonas* sp.). The latter association was found in ice-covered lakes as well, suggesting a common link between this chrysophyte taxon and its fungal peer (Rojas-Jimenez et al. 2017). Most interactions between chytrids and algae involved multiple partners which suggests that these potential parasitic interactions work in multifaceted ways to impact host distribution and abundance. However, it is important to note that these potential pathogens might have saprophytic lifestyle, too, participating in the decomposition processes of the host cells. In that scenario, our prediction of the ecological role of the associated partners can be greatly challenged (Egan & Gardiner 2016).

Previous studies suggested the importance of Cryptomycota (e.g., *Rozella*) species in the regulation of population size of parasitic zoosporic fungi in lakes (Gleason et al. 2012). We found, in contrast to our initial assumption, no sign of Cryptomycota OTU which may have been acted as hyper-parasites of parasitic chytrids (i.e., chytrid showing strong associations with algal OTUs). Although some strong links were found between Cryptomycota OTUs and Chytridiomycota OTUs, these chytrids were hypothetically saprotrophs as they lacked clear relationship with any algae. Hence, we cannot rule out the possibility that these Cryptomycota–Chytridiomycota links might indicate mere co-distributions or other biotic interactions. Nevertheless, detected associations between members of Cryptomycota (unassigned) and Dinoflagellata suggest that this fungal group might have a relevant role in the regulation of algal populations (e.g., *Dinophysis* and *Prorocentrum*) that can cause blooms in the Baltic Sea (Gisselson et al. 2002, Hajdu et al. 2005) without being targets for chytrids (Gleason et al. 2015). This, in turn, emphasizes that Cryptomycota may have as important role as chytrids in aquatic food webs, and reinforce the existence of other than chytrid-based mycoloops in aquatic ecosystems (Kagami et al. 2014).

### Conclusions and future perspectives

This study enhances our knowledge of fungal diversity in coastal marine habitats, elucidate their spatiotemporal variation, and presents biotic (i.e., parasitic) interactions with planktonic algae. We believe that the joint application of distribution model and network approaches represents the advantage to infer a more detailed picture of metacommunity assembly with the inclusion of species covariation (attributed to biotic interactions), and can be used to support conclusions drawn from network results (e.g., presence of biotic interactions between Cryptomycota and chytrids).

Most previous studies investigating fungal interactions have mainly been restricted to studies in laboratory settings. Whilst those studies deepen our knowledge in biotic interactions and their mechanisms, they barely provide information about how these interactions actually impact the distribution of mycoplankton communities in nature. Together, our findings emphasize that the contribution of biotic associations to fungal metacommunity assembly are important to consider in future studies as it helps us to improve predictions of species distributions in aquatic ecosystems. Furthermore, identifying biotic relationships that affect the distributions of members of mycoplankton could be useful for plankton ecology, through habitat management to promote species which control algal blooms and facilitate nutrient transfer to upper trophic levels. Nevertheless, it is important to note that the putative interactions we present in this study may not merely capture true biotic interactions involving parasitic relationships but could reflect the effect of relevant missing predictors (Wilkinson et al. 2021).

Investigations targeting aquatic fungi, with a special focus on “dark matter fungi”, have a great potential. Therefore, future works should extend chytrid-centered studies on other fungal groups (e.g., Cryptomycota) and consider studying their ecological roles and community assembly in a wider selection of aquatic ecosystems. We also imagine future field studies as being the basis of laboratory experiments wherein putative biotic interactions could be further assessed, moreover, the ecological role of mycoplankton members could be investigated using more directed (e.g., single-cell genomics and meta-omics) approaches (Laundon & Cunliffe 2021). Taken together, we emphasize the need to go a step beyond culture-based studies and approach aquatic fungi from the (meta)community level perspective in future studies.

## Acknowledgements

We are especially thankful to Juanjo Rodriguez Serrano who made it possible for us to set-up and perform sequencing on the Nanopore Mk1C, and Tomasz Dobrzycki at ONT for giving advices during the sequencing. We are grateful for the support of Sonia Brugel and Umeå Marine Science Centre for collecting samples and performing physicochemical measurements, Kristoffer Sahlin for bioinformatic support, and Maximilian Pichler for his help and advice for the joint species distribution modeling. Funding for this project was provided by Umeå University and the Swedish strategic marine research programme EcoChange (Swedish Research Council Formas). This research was conducted using the resources of High-Performance Computing Center North (HPC2N). The computations and data handling were enabled by resources provided by the Swedish National Infrastructure for Computing (SNIC), partially funded by the Swedish Research Council through grant agreement no. 2018-05973.

## Data availability

The OTU tables, their taxonomic classifications, and the environmental data related to the samples, as well as the model R data file are deposited in Open Science Framework (OSF) (DOI: 10.17605/OSF.IO/764MU).

## Author contributions

AA, UCG, JW conceived the study and obtained funding for this research. KE performed sample processing and provided samples for this study. MV performed further sample processing prior to sequencing, developed the research questions, analysed the data and drafted the manuscript. All authors contributed substantially to writing and revisions.

