## Supplementary material for "Co-occurrences enhance our understanding of aquatic fungal metacommunity assembly and reveal potential host–parasite interactions"

**Amplification details:**

PCR was performed in 40 µL reactions with 1.5 µL PrimeStar GXL polymerase (#R050A; Takara), 12 pmol of barcoded primer (see list of barcodes in Supplementary Table S1), 1 mM dNTPs, 8 µL 5× PrimeSTAR GXL Buffer and 2 µL of DNA template (0.9–49.6 ng/µL). Amplifications were done on a Bio-Rad T100 Thermal Cycler (Bio-Rad Laboratories) with an initial denaturation at 98 °C for 1 min, then 36 cycles at 98 °C for 10 s, annealing at 55 (or 60) °C for 15 sec and elongation at 68 °C for 4 min.

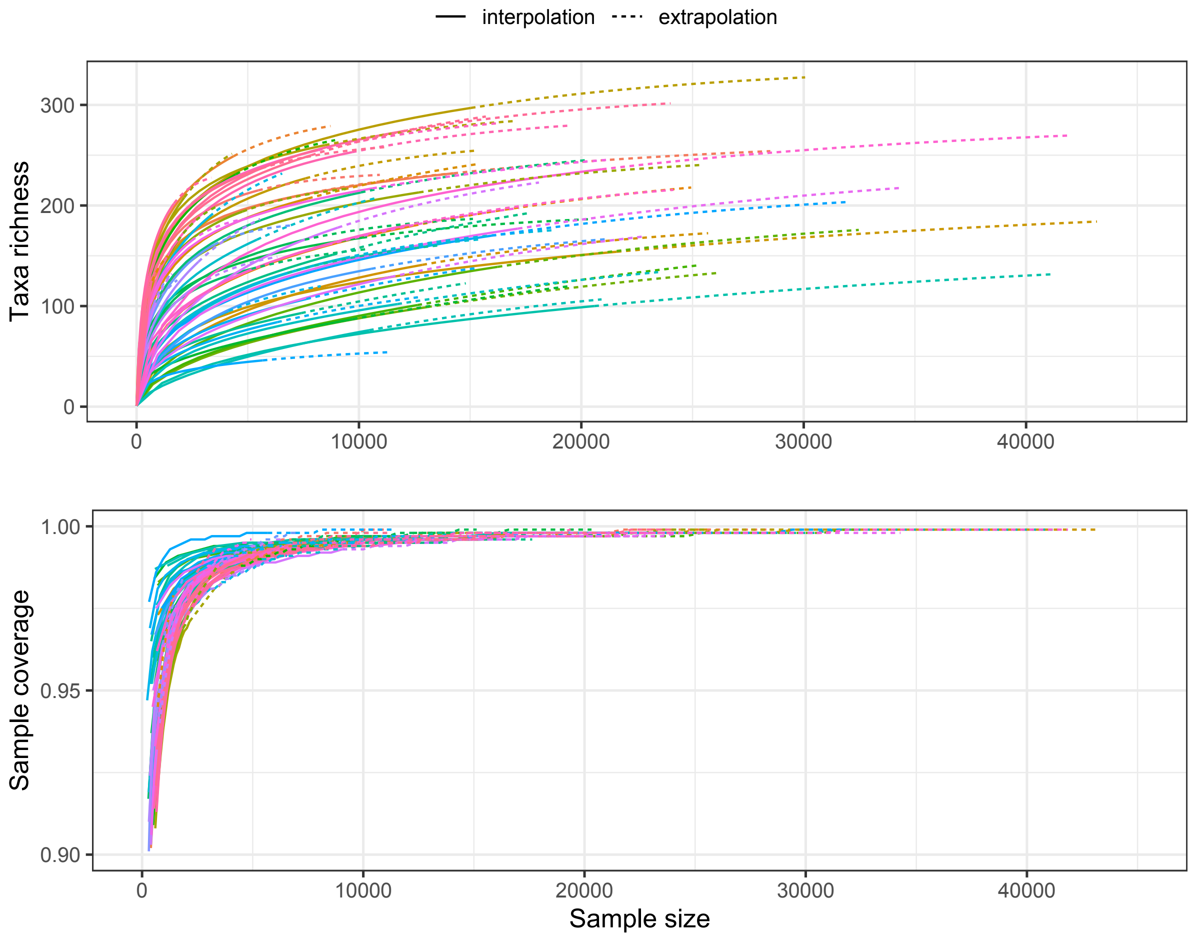


**Figure S1.** Diversity estimates and sample completeness (coverage) curves.

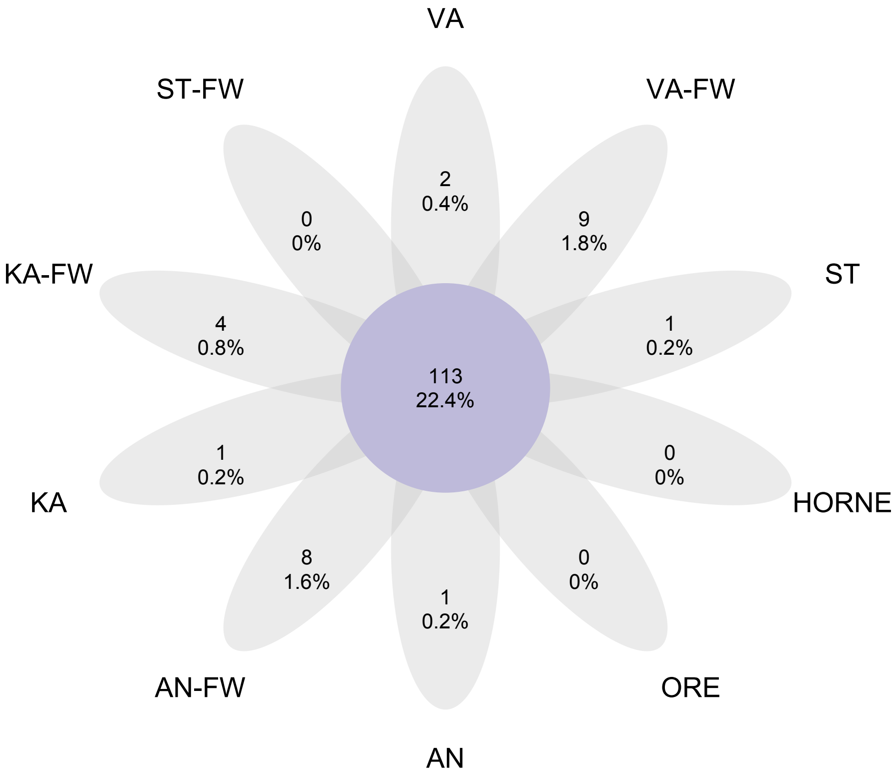


**Figure S2.** Venn diagram representing the unique and shared fungal taxa in the sampled bays and their freshwater inlets ('-FW' tag), or in the offshore sites (HORNE, ORE). For code descriptions, see Figure 1. in the main text.

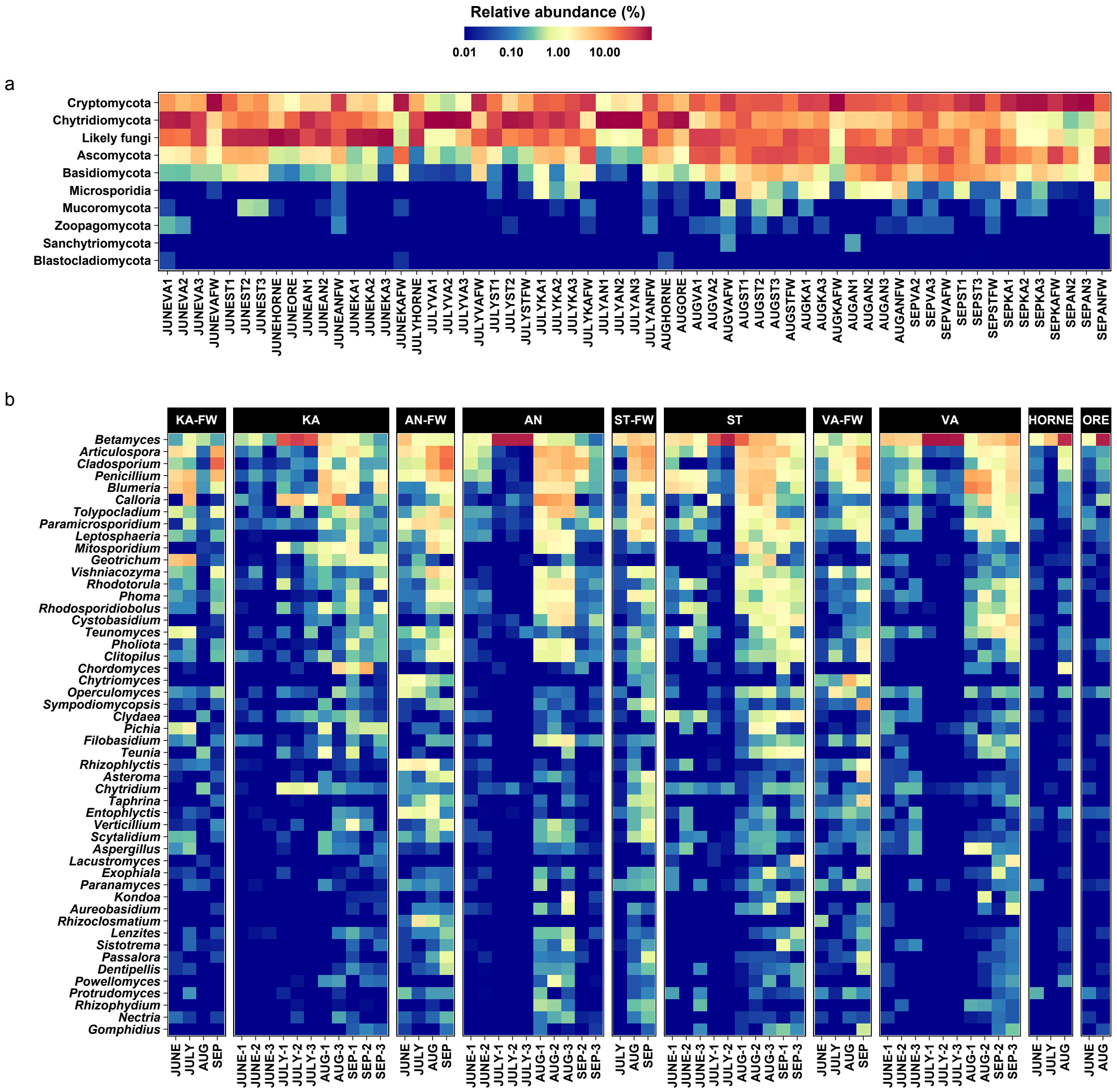


**Figure S3:** Heatmaps showing the temporal changes of fungal communities at (a) phylum and (b) genus level. Note that only the top 50 OTUs assigned (based on BLAST search) down to genus level are listed.

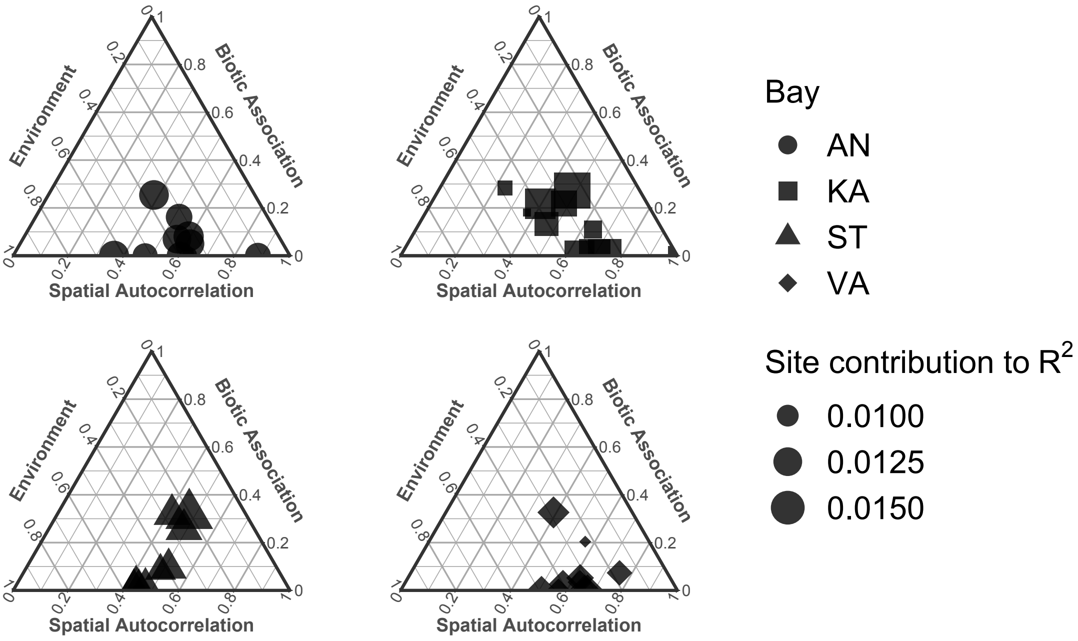


**Figure S4.** Site-specific internal metacommunity structure. The ternary plots describe the contributions of space, environment and biotic associations to the metacommunity level properties among the four bays. The size of the symbols refers to the variation explained by the model ($R^2$) for each site.

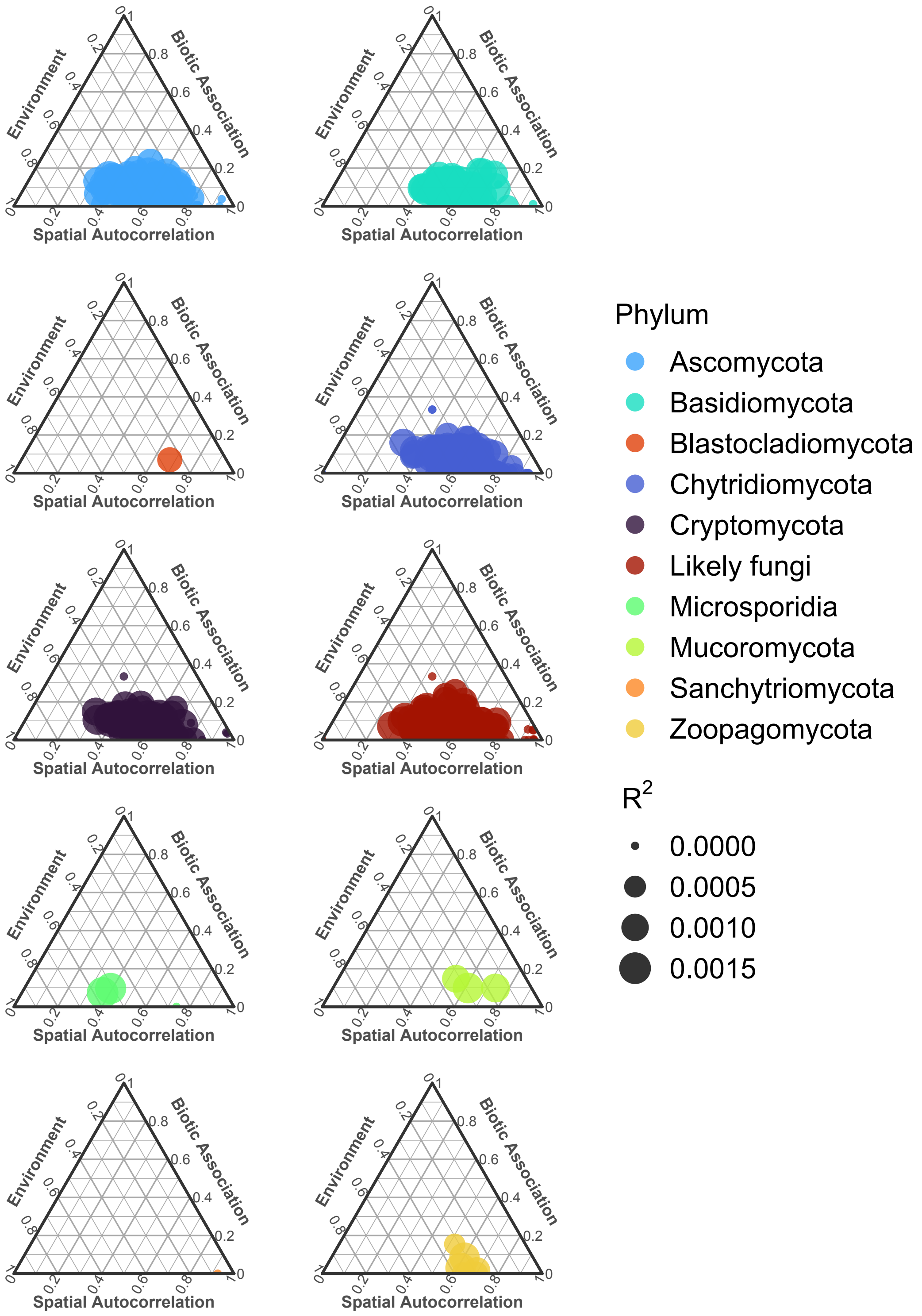


**Figure S5.** Taxa-specific internal metacommunity structure. The ternary plots display the relative influence of environmental conditions, space and biotic associations on the distribution of each fungal taxon (OTU). The size of the symbols refers to the variation explained ($R^2$) by the model for each OTU.

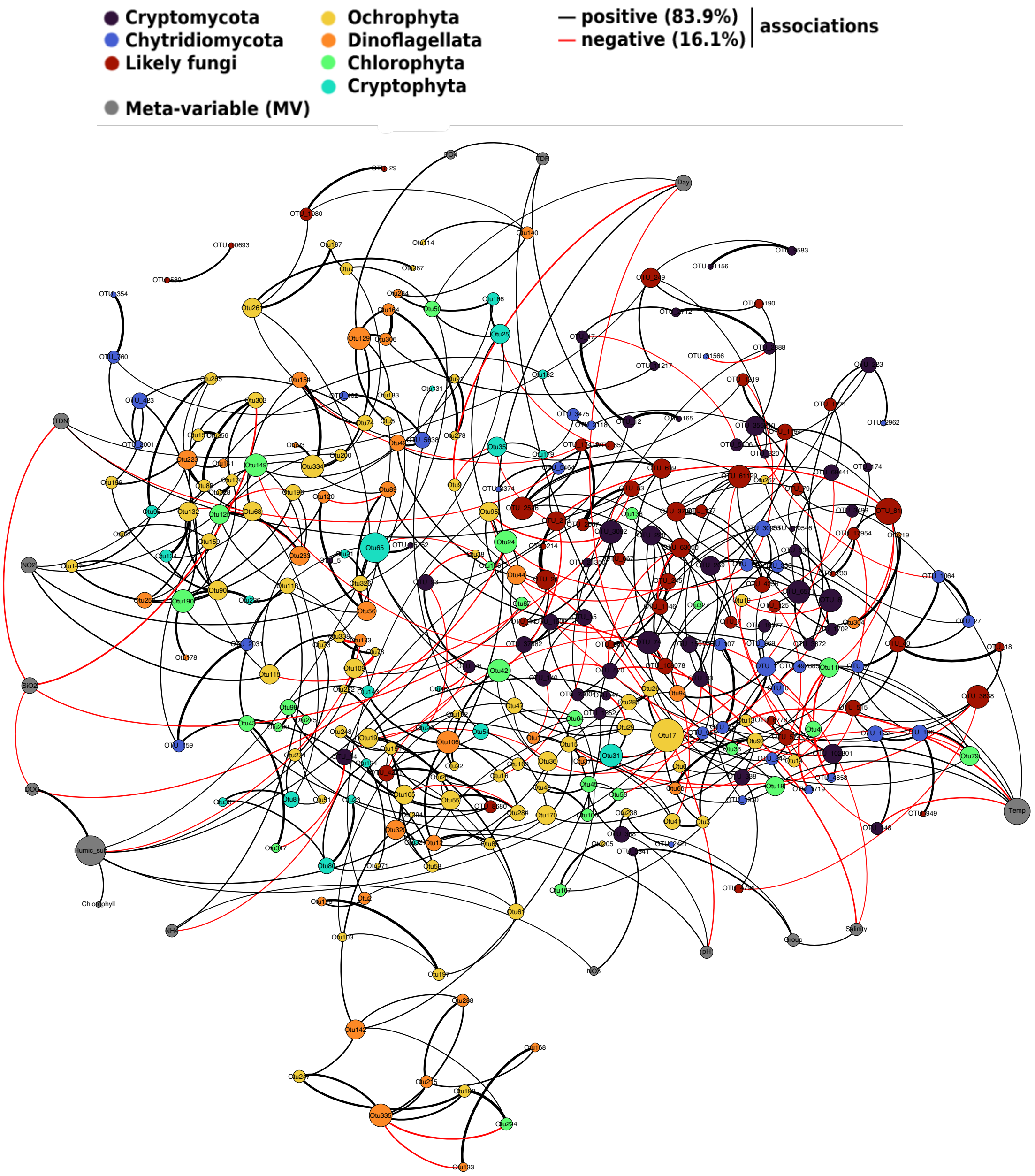


**Figure S6.** Prediction of fungi–algae interactions with FlashWeave (alpha < 0.05). The size of nodes (OTUs or meta-variables; n = 303) are proportional to the number of predicted interactions (degree). Edge width refers to the strength of correlations (weight cutoff > |0.3|; n = 498) and coloured by association type (positive or negative). 'Likely fungi' refers to OTUs that were assigned as fungi (based on BLAST search) but without matches to any phylum.
